# Artemin and its cognate receptor, GFRα3, play a function role in osteoarthritis pain

**DOI:** 10.1101/2020.11.04.368373

**Authors:** Laura Minnema, Santosh K. Mishra, B Duncan X Lascelles

**Author notes:** Corresponding authors: SK Mishra; BDX Lascelles.

## Abstract

Osteoarthritis associated pain (OA-pain) is a significant global problem. OA-pain limits limb use and mobility, and is associated with widespread sensitivity. Therapeutic options are limited, and the ones that are available are often associated with side or adverse effects. The lack of therapeutic options is partly due to a lack of understanding of clinically relevant underlying neural mechanisms of OA-pain. In previous work in naturally occurring OA-pain in dogs, we identified potential signaling molecules (artemin/GFRα3) that were upregulated. Here, we use multiple approaches including knockout mice, immunological suppression in a mouse model of OA, and clinically relevant measures of sensitivity and limb use to explore the functional role of artemin/GFRα3 signaling in OA-pain. We found the monoiodoacetate (MIA)-induced OA model in mice is associated with decreased limb use and hypersensitivity. GFRα3 expression is increased in sensory neurons. Exogenous artemin induces heat, cold and mechanical hypersensitivity, and anti-artemin monoclonal antibody administration reverses this hypersensitivity and restores limb use in mice with MIA-induced OA pain. Our results provide a molecular basis of arthritis pain linked with artemin/GFRα3 signaling and indicate that further work is warranted to investigate the neuronal plasticity and the pathways that drive pain in OA.

## Introduction

While the term ‘arthritis’ encompasses around 100 different types of joint disease, osteoarthritis (OA) is one of the most common forms of degenerative joint disease of humans and companion animals. The prevalence of OA is around 35.5 % worldwide (1) and similar for companion dogs and cats (2-4).

This joint disease is multifactorial with multiple factors contributing to the disease and whether or not pain is associated with OA, but the pain is the overriding complaint and cause of disability. Ongoing, chronic pain affects multiple dimensions, and causes deterioration in the musculoskeletal system, results in algoplasticity and increased sensitivity, as well as having negative effects on the affective system, cognitive function and relationships (5). There are few effective treatments for OA available, and these are often associated with severe side effects and safety concerns. Treatments include corticosteroids, non-steroidal anti-inflammatory drugs (NSAIDs), and the soon to be approved anti-nerve growth factor (NGF) monoclonal antibodies (6,7). Corticosteroids can be effective at relieving joint pain, but due to the need for intra-articular injections every 1-4 weeks they are mainly used as a short-term solution (8), and repeat injections raise concerns about systemic exposure and effects on joint cartilage. NSAIDs can be associated with severe side effects, including gastrointestinal bleeding and increased risk of heart attack or stroke (9). Adverse effects of anti-NGF include transient paresthesia and edema, rapidly progressive OA (∼1.5-3.0%), and, in a small number of patients treated with both anti-NGF and NSAIDs, osteonecrosis. The adverse effects of anti-NGF are dose-dependent, with lower doses being safer, but lower doses are also less effective for pain-relief (10). In brief, problems associated with these above therapeutics potentially limit the treatment for OA-pain. Clearly, a better understanding of the molecular underpinnings of OA pain is needed in order to develop novel and safe therapeutics.

Recently, we found an elevated serum level of artemin in humans and dogs with naturally occurring OA (11). Further, we found significantly increased GFRα3 and transient receptor potential vanilloid (TRPV) subfamily-1 receptor expression in dogs DRG serving osteoarthritic joints compared to healthy dogs. Additionally, we showed that synovial fluid artemin concentrations were associated with joint pain. Overall, results from our work in the naturally occurring OA dog model suggest a possible role of artemin and its receptor GFRα3 in OA-pain. To examine the functional role of artemin/GFRα3 signaling in OA-pain, here we used a mouse model of OA based on the intra-articular injection of mono-iodoacetate [MIA], a glyceraldehyde-3-phosphatase dehydrogenase inhibitor. The MIA model has been extensively used in the rat and well-characterized for pain phenotypes since the first descriptions in 1985 (12). The MIA model has been shown to have characteristics of human OA pathogenesis and pain (13). Less work has been performed with the MIA model in mice, and the algoplastic changes in heat, cold, and mechanical sensitivity have not been fully described. Not only are these ‘standard’ measures of sensitivity associated with pain, but human OA patients demonstrate changes in response to these stimuli. (14)

After confirming the development of heat, cold, and mechanical sensitivity in the mouse MIA model, we further explored the role of GFRα3 and TRPV1 ion channels in OA-pain. Finally, we showed artemin-induced mechanical and thermal pain sensitivity were attenuated in mice administered with an anti-artemin monoclonal antibody. In summary, our results provide a significant role of artemin/GFRα3 signaling in OA-pain.

## Results

### MIA dose-determination work

We tested three doses of MIA (0.1, 0.5, or 1 mg dissolved in 10 μl of saline) to determine an appropriate dose for subsequent experiments in mice. Sensitivity to heat, cold, and mechanical stimuli was evaluated in mice that received injections of saline (control), 0.1 mg, 0.5 mg, or 1 mg of MIA. For all doses, all mice survived to Day 28. All mice that received MIA showed hypersensitivity compared to control mice. Mice that received 1 mg of MIA showed the greatest degree of hypersensitivity to heat, cold, and mechanical stimuli through to Day 28. With only two mice per group for this preliminary work, statistical power was limited, but there was a significant difference between control mice and 1 mg MIA at day 28 for all three behavioral assays [Supplementary Figure 1 (two-tailed t-test assuming unequal variances; von-Frey p<0.05; Hargreaves p<0.05; cold p<0.01)]. For all subsequent work, we used 1 mg MIA in a 10 μL volume (1 mg/10 μL).

### MIA-induced OA pain is associated with mechanical, heat and cold hypersensitivity

Next, we evaluated mice for mechanical, heat, and cold hypersensitivity after intra-articular injection with either MIA (1 mg/10 μL) or saline (10 μL). Mice were tested at 1, 2, 3, 7, 14, 21, and 28 days following injection of MIA. Sensitivity of the contralateral limb of the MIA-injected mice did not differ from the contralateral or ipsilateral limbs of the saline-injected mice (data not shown). The ipsilateral (injected) limb of MIA mice was hypersensitive to mechanical, hot, and cold stimuli at all time points, compared to the ipsilateral limb of the saline-injected controls (Figure 1).

**Figure 1.**
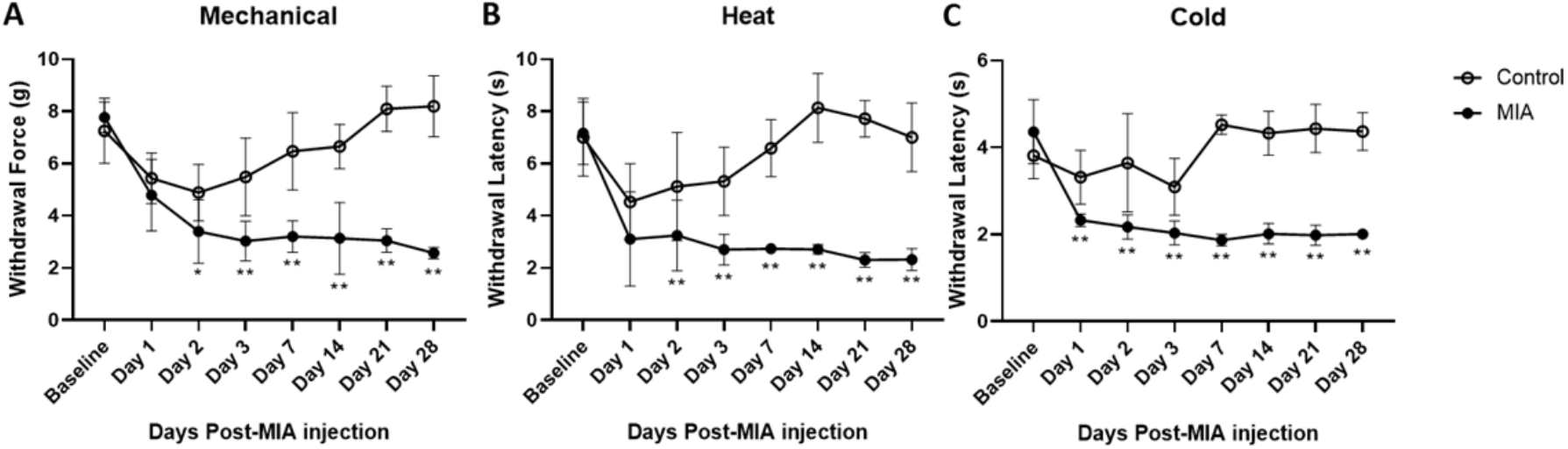
Hypersensitivity to mechanical, thermal, and cold stimuli after MIA induction. Mice were tested for hypersensitivity to mechanical (A), heat (B), and cold (C) stimuli using the Von Frey, Hargreaves, and crushed dry ice assays over 28 days following MIA injection. MIA (n=5) and control (n=6). Data are represented as mean ± SD. Significance based on a repeated measures ANOVA with post-hoc multiple comparison adjustments (p<0.05).

### MIA-induced OA pain reduces bodyweight distribution to the painful limb

A separate cohort of mice received an intra-articular injection of either MIA (1 mg/10 μL) or saline (10 μL). A static horizontal incapacitance meter (SHIM) (15) was used to assess the percent of hindlimb supported body weight placed on each hindlimb. Mice placed significantly less weight on the injected limb compared to the contralateral limb starting at Day 2 and continuing to Day 28 (Figure 2a). Figure 2b shows the asymmetry in weight bearing on the limbs.

**Figure 2.**
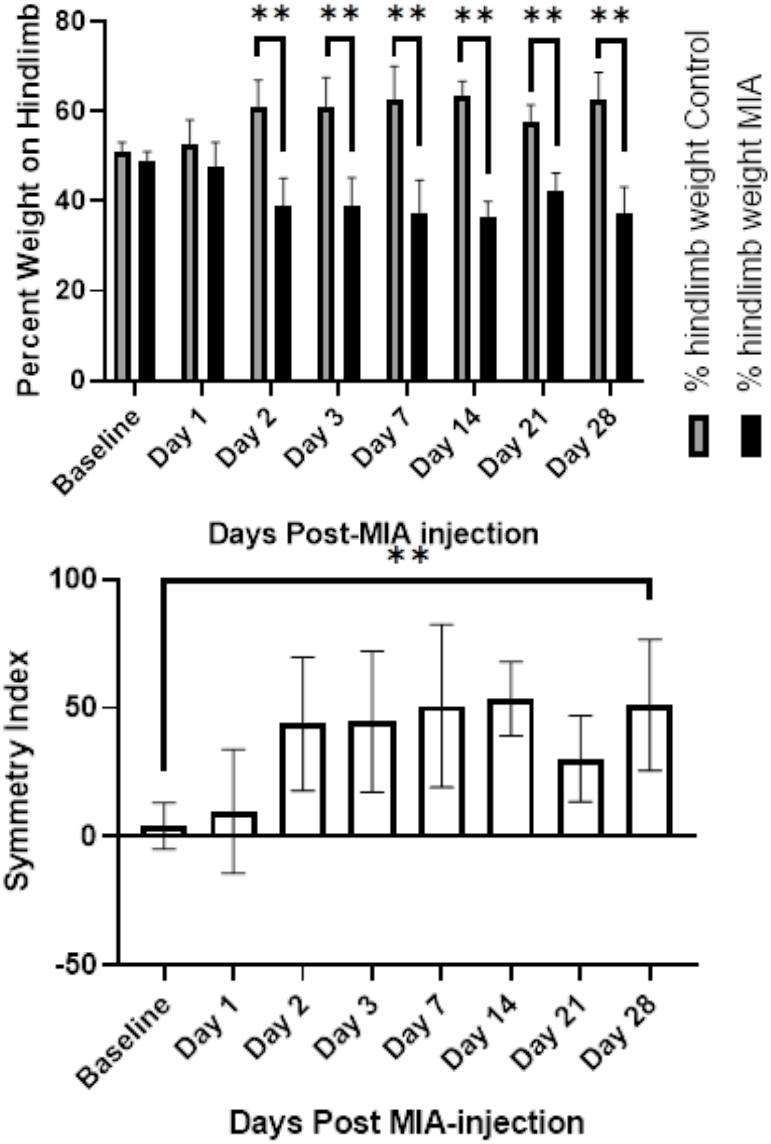
Static weight-bearing after MIA induction. A) Standing bodyweight distribution to the hindlimbs was measured over 28 days following MIA OA induction in mice (n=8), and percentage of total weight placed on the hindlimbs that was born on the ipsilateral and contralateral limbs is plotted. The MIA-injected hindlimb placed significantly less weight on the ipsilateral (MIA) limb compared to the control contralateral limb from day 2 onward (ANOVA with repeated measures post-hoc testing; ** = p<0.01). Data are represented as mean ± SD. B) Static weight-bearing data represented as a symmetry index. Greater asymmetry in weight bearing was observed after MIA induction compared to baseline, following the same pattern of being significantly different from baseline starting at day 2 after MIA-induction (p<0.05).

### Role of TRPV1 in MIA-induced OA hypersensitivity

TRP channels play a role in encoding noxious stimuli. To determine the contribution of TRPV1 in MIA-induced OA pain, we use TRPV1 knock out (KO) mice and compared responses to control littermates. TRPV1 KO mice had an increased withdrawal latency compared to wild-type mice at baseline; i.e., the TRPV1 KO mice were less sensitive to heat, as expected. However, TRPV1 KO status did not protect mice from heat hypersensitivity induced by MIA (Figure 3B). Further, TRPV1 KO status had no effect on mechanical and cold hypersensitivity induced by MIA (Figure 3A and 3C).

**Figure 3.**
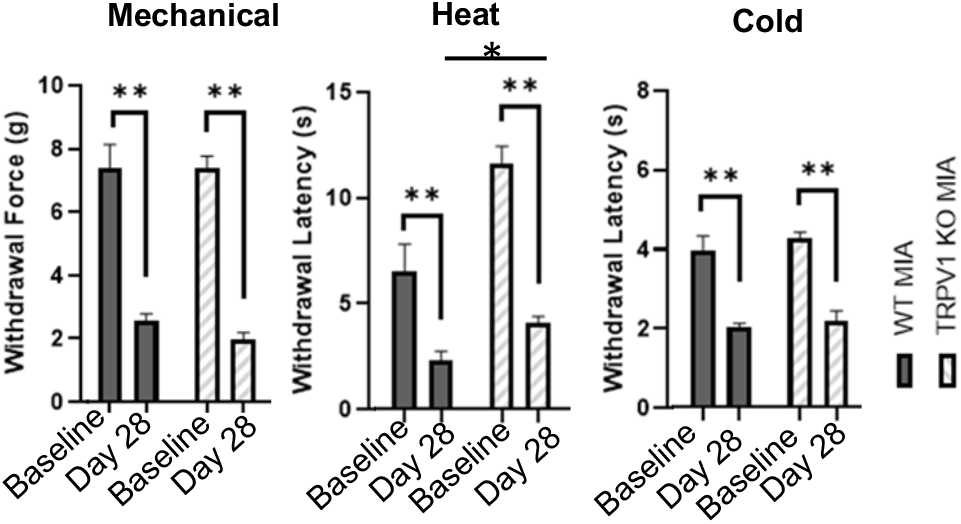
MIA induced hypersensitivity in TRPV1 KO mice. Mechanical (A), heat (B), and cold (C) sensitivity at baseline and at day 28 following MIA induction of OA in WT and TRPV1 KO mice. (A) Both WT and KO mice given MIA showed a significant reduction in withdrawal force (g) at day 28 (p<0.01; p<0.01). (B) TRPV1 KO mice had significantly longer withdrawal latencies (less sensitive to heat) at baseline than WT mice (p<0.01). At day 28 TRPV1 KO MIA mice had less hypersensitivity to heat than WT MIA mice (p<0.01). (C) TRPV1 KO mice were no different from WT mice (p>0.05), and both WT and TRPV1 KO mice showed significant cold hypersensitivity at day 28. Data are represented as mean ± SD, n=6 mice each group.

### GFRα3 receptor expression is increased in MIA OA pain

Recently, we showed an increase in the expression of GFRα3 in the DRG of naturally occurring osteoarthritic dogs (11). Here, we sought to determine whether the same change held for the MIA-model of OA in mice. Using immunofluorescence staining, we found that GFRα3 was expressed in about 40% of neurons of DRG serving MIA-injected joints compared to ∼20% of neurons serving the contralateral joints and DRG serving joints that received saline (Figure 4A, 4B, and 4C). Using double IHC we found that the majority of the GFRα3 expression occurs in TRPV1 expressing neurons, although in the MIA state there is some de-novo expression in non-TRPV1 expressing neurons (supplementary figure 2).

**Figure 4.**
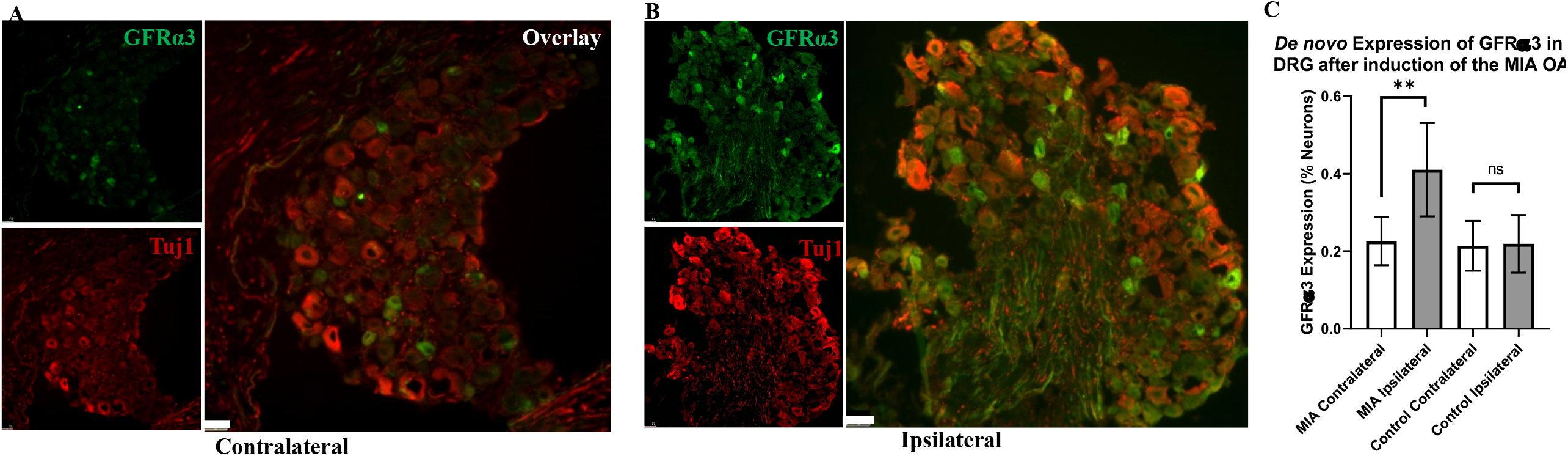
*De novo* expression of GFRα3. (A) and (B) shows representative contralateral and ipsilateral double immunostaining on mouse DRG sectioned stained with Tuj1 in red and GFRα3 in green. (C) Quantification of GFRα3 normalized against a pan neuronal marker (Tuj1) revealed an increase in GFRα3 in mice treated with MIA. Around 10 sections from each side were counted for number of neurons (Tuj1) and number of neurons expressing GFRα3, n= 3 sham and 3 MIA mice (p=0.016963). We found no difference between saline controls (ipsilateral and contralateral) and contralateral MIA DRG (p=0.84 and p=0.90).

### Artemin, a ligand of GFRα3 receptor, plays a role in thermal and mechanical hypersensitivity

OA is associated with thermal and mechanical hypersensitivity. In previous work we have determined that artemin, the ligand for GFRα3, is increased in synovial fluid and serum of dogs, and serum of humans, with OA. We therefore sought to determine if artemin is associated with thermal and mechanical hypersensitivity. We found that injection of artemin into the ventral surface of the paw was associated with significantly increased sensitivity to heat, cold, and mechanical stimuli, as shown in Figure 5. Thermal hypersensitivity to heat (Figure 5A) and cold (Figure 5B) lasted from 1 to 4 hours post-injection while mechanical hypersensitivity (Figure 5C) had a slightly delayed onset, lasting from 2 to 6 hours post-injection. All mice returned to baseline by 24 hours post-injection.

**Figure 5.**
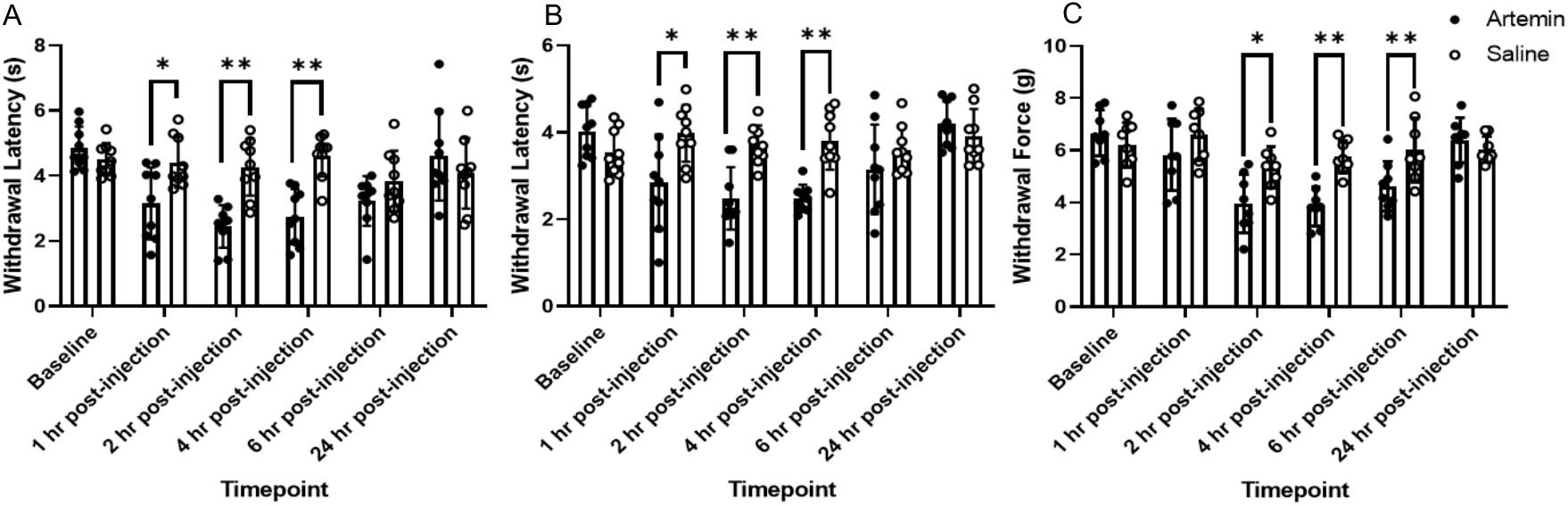
Artemin-induced hypersensitivity to heat, cold, and mechanical stimuli. Mice that received artemin had significantly reduced withdrawal latencies to heat and cold (A,B) compared to mice that received saline at 1 hr (p<0.05), 2 hr (p<0.01), and 4 hr (p<0.01) post injection (multi-comparison t-test; n=9 per group). Mechanical withdrawal force (C) was significantly decreased at 2 hr (p<0.05), 4 hr (p<0.01), and 6 hr (p<0.01) post-injection with artemin (multi-comparison t-test; n=8 per group). Data are represented as mean ± SD.

### Blocking signaling between artemin and GFRα3 attenuates thermal and mechanical hypersensitivity in MIA OA pain

MIA injected into the right stifle joint was used to induce OA as described earlier. Mice were tested weekly for 28 days to confirm hypersensitivity. Anti-artemin monoclonal antibody [25 mg/100 μL; intraperitoneal (i.p.)] was administered on day 28. Mice that received the antibody demonstrated reduced hypersensitivity compared to controls between 2- and 4-hours post-injection (Figure 6) for mechanical and cold, and out to 6 hours post-administration for heat. All mice were back to pre-injection response levels by 24 hours of post-anti-artemin injection.

**Figure 6.**
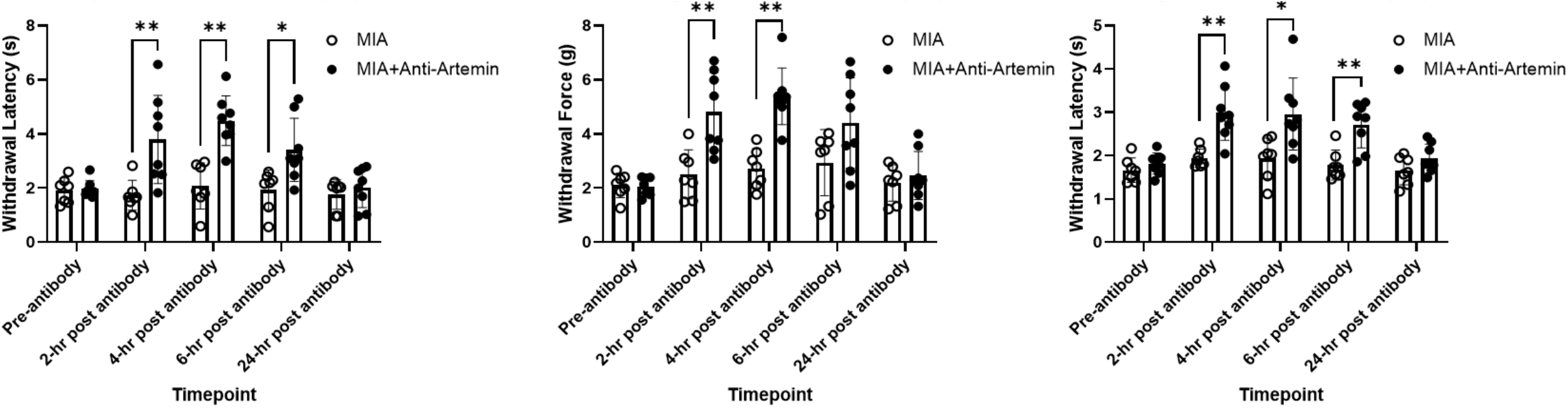
An anti-artemin antibody in alleviating hyperalgesia in the MIA model of OA. Anti-artemin alleviates MIA-associated hypersensitivity for heat (A), mechanical (B), and cold (C). Anti-artemin was effective in reversing thermal hypersensitivity to hot and cold stimuli at 2, 4, and 6 hr after antibody administration and at 2 and 4 hr post antibody for mechanical hypersensitivity (t-test; p<0.05). Data are represented as mean ± SD.

## Discussion

Here, using a mouse model of OA, and clinically relevant measures of sensitivity and limb use, we explored the functional role of artemin/GFRα3 signaling in OA-pain. We found the monoidoacetate (MIA)-induced OA model in mice is associated with decreased limb use and hypersensitivity. TRPV1 KO mice that have OA induced using MIA have partial reduction in pain but still show residual heat, mechanical, and cold sensitivity, suggesting signaling pathways other than TRPV1 are responsible for heat hypersensitivity in OA-pain. We found that in MIA-induced OA-pain, GFRα3 expression is increased in sensory neurons, and expression extends outside of the baseline expression in TRPV1-expressing neurons. We found that exogenous artemin induces heat, cold, and mechanical hypersensitivity, and anti-artemin monoclonal antibody administration reverses MIA-induced evoked pain hypersensitivity. Overall, our results indicate artemin/GFRα3 plays a functional role in arthritis pain.

### Mouse MIA model of OA-pain

Mechanisms of OA-pain have been widely studied in rat models employing intra-articular injection of chondrocyte glycolytic inhibitor mono-iodoacetate (MIA) because of the relative ease of stifle joint injection in rats, and the relatively rapid onset of joint destruction. However, in mice, only a few groups have reported the use of MIA and demonstrated pain phenotypes (16-21). Additionally, the doses used have varied widely (22)We confirmed an appropriate dose (1 mg) and showed that in mice with MIA-induced OA, there is increased heat, cold, and mechanical hypersensitivity compared to controls. These changes in the sensitivities mirror what is seen in humans with OA-pain (23) and dogs with naturally occurring OA-pain (24,25). Approximately 70% of human patients with knee OA experience hypersensitivity to at least one modality – mechanical, hot, or cold (26,27). In a separate experiment we measured bodyweight distribution using a static horizontal incapacitance meter and found a reduction in limb use suggesting ongoing joint pain. Human knee osteoarthritis patients also display weight-bearing asymmetry while at rest with less weight born on the affected limb, (28-30) again demonstrating the relevance of the murine MIA model of OA-pain. To best of our knowledge, no other laboratories have confirmed the combination of heat, cold, and mechanical hypersensitivity and decreased limb use in the mouse MIA model.

### Role of TRPV1 ion channel in OA-pain

The TRPV1 receptor has been widely studied for its role in the transduction of stimuli and the generation of pain induced by noxious heat (18,31). Recently, the role of TRPV1 in pain and sensitivity associated with OA-pain has been reported across various models and species. In one study in rats, a TRPV1 antagonist blocked the development of thermal (heat) hypersensitivity, but not weight-bearing deficits or ongoing pain (32). In a complete Freund’s adjuvant (CFA) stifle OA model in mice, the TRPV1 antagonist, A-425619, reversed the decrease in digging behavior induced by CFA injection. There are no reports of the role of TRPV1 in the murine MIA model of OA-pain. TRPV1 antagonism has only shown mild therapeutic effects in humans (33) and a high incidence of adverse events; and in dogs with naturally occurring OA-pain; a TRPV1 antagonist was not effective. (34) In our study, we used TRPV1 KO mice to examine if TRPV1 is involved in MIA-induced heat, cold, and mechanical hypersensitivity. Not to our surprise, we found the loss of function of TRPV1 was partially protective for heat sensitivity induced by MIA at Day 28 but TRPV1 KO had no effect on mechanical and cold hypersensitivity. The role of TRPV1 in detecting heat but not mechanical sensation is consistent with previous reports (18,35). Collectively, the literature and our results indicate little evidence of an analgesic effect of blocking or removing the influence of the TRPV1 receptor, despite its known role in pain processes. This suggests that other signaling systems are involved in OA-pain, and upstream signaling mechanisms may be more effective therapeutic targets.

### MIA-induced de novo expression of GFRα3 in murine DRGs

Recently, we identified an increase in the expression of GFRα3 at both protein and RNA level in the naturally occurring canine model of OA-pain, suggesting this change in the GFRα3 protein is due to underlying disease conditions. The GFRα3 receptor acts upstream of TRP receptors. The role of GFRα3 in pain has been partially established through work showing GFRα3 signaling in cold pain (36), bladder pain (37), and inflammatory bone pain (38). However, the role of GFRα3 signaling in arthritis pain has not been investigated to date. Interestingly, our data suggests a *de novo* expression of GFRα3 in the DRG, doubling the number of neurons that are expressing the GFRα3 receptor, perhaps indicating a change in neuronal plasticity of sensory neurons. Increased GFRα3 expression might be a mechanism that contributes to widespread pain sensitivity through TRP channel activation in OA, but this needs to be investigated in future work.

### Role of artemin/ GFRα3 signaling in OA-pain

OA is a degenerative disease, and in the process of tissue degeneration, various mediators are released. Based on our data from naturally occurring osteoarthritic dogs, and confirmed in a small number of human samples, we found artemin was increased in the serum in association with OA-pain. (11) Data from our present study clearly show that a single local injection of artemin into a mouse’s paw can induce hypersensitivity to thermal, cold, and mechanical stimuli as well as decrease weight bearing – all features of human OA patients. These data are in agreement with a report suggested repeated application of artemin increase mechanical and heat hypersensitivity (39). Conversely, in another study, investigators found no change in mechanical response post artemin injection (36). Interestingly, cold hypersensitivity in response to artemin injection was consistently seen (36). The studies by Lippoldt et al., used both sexes of mice however, we used only male mice in our study, and this may explain some of the differences for heat and mechanical hypersensitivity. We used only male mice in our study to avoid the impact of the estrous cycle on pain sensitivity (40) in these early experiments, but future experiments should use both sexes.

The expression of GFRα3 in naïve DRG and its co-expression and interaction with nociceptive channels is known, but its role in chronic pain state is yet to be explored. Reports have shown that artemin caused an increase in the mRNA for GFRα3, tropomyosin receptor kinase A (TrkA), TRPV1 and TRPA1 (41) and an anti-artemin antibody was shown to block upregulation of TRPA1 (37) and knockdown of GFRα3 with siRNA blocked the upregulation of TRPV1 in a nerve injury model (42). In the results we report here, we show that blocking the signaling between artemin/GFRα3 by antibody sequestration of artemin in the established mouse MIA model of OA-pain method inhibits heat, cold, and mechanical sensitivity. The difference between the effect of the TRPV1 KO status and mAb anti-artemin suggests that blocking artemin/GFRα3 signaling may have more analgesic potential than targeting individual TRP receptors because it is upstream of these nociceptive channels.

Overall, we present the first evidence of a potential functional role of artemin/GFRα3 in chronic OA-pain. However, there is much to understand about the potential role of artemin, including the mechanisms leading to artemin release and which cells types in the joints are responsible for artemin release; whether artemin is involved in the induction and/or maintenance of OA-pain; and whether artemin acts through only GFRα3, or other receptors (such as NCAM) are involved. Also, the degree to which artemin may drive the ultimate experience of OA pain needs to be fully elucidated. Although we have found that artemin’s cognate receptor, GFRα3, was upregulated in MIA induced OA-pain, paralleling what we found in the naturally occurring OA model in the dog, work needs to be done to determine the contribution of GFRα3 upregulation and/or activation in OA pain, and further, the downstream targets and signaling mechanisms need to be defined.

## Materials and Methods

All experiments were performed under institutional animal care and use committee approval (IACUC # 19-047B).

### Mice

In all experiments, across all groups, aged-matched, 4-week-old, adult male C57BL6 mice (Jackson Labs) were used. The average mouse weight was 25 grams and mice were housed in groups of four and kept on a 12-hr light-dark cycle. TRPV1 knockout (KO; Jackson Labs-Stock No: 003770) mice were on a C57BL6 background.

### Preparation and injection of MIA

Intra-articular injection of MIA was performed according to the method published by Pitcher et al. (2016). Following pilot dose-determination experiments, MIA (Sigma-Aldrich, I2512) was dissolved in sterile saline and 10 μl containing 1mg MIA was injected intra-articularly using a zero-dead space syringe with a 30G needle (Hamilton). Control mice received 10 μl of sterile saline intra-articularly. Right stifles were used for all injections. Mice were anesthetized for the injection using a nose-cone delivering 5% isoflurane carried in oxygen. The injection site (right stifle) was cleaned with surgical scrub (70% Ethanol) and iodine solution three times prior to the injection. To reduce inflammation and abrasions, we did not shave the injection area, deviating from the published protocol (Pitcher). Instead, the surgical scrub was used to flatten and part the fur for visualization of the injection site. The joint space was identified by flexing the leg and using a transversely applied needle to identify the distal patella ligament and underlying joint space (43). Mice were under anesthesia for no longer than 5 minutes.

### Behavior

All behavioral assays were performed by the same researcher (LM) to maintain consistency, and LM was blinded to treatment groups to minimize bias. Behavioral assays were conducted at the same time of day (afternoon) for each time point. Mice were acclimated to the testing environment and each piece of equipment for 5 minutes before each testing. At each time point, behavioral assays were repeated five times with five minutes between each measurement.

#### Evoked pain behavior

Mechanical sensitivity was measured using the Ugo Basile Dynamic Plantar Aesthesiometer, referred to as the electronic Von Frey. A mechanical stimulus is delivered to the hind paw and the force at which mouse withdraw its paw was recorded. Heat sensitivity was tested using the plantar assay (or Hargreaves apparatus, Ugo Basile). Mice were placed in testing chambers on a glass plate. An infrared light source was focused on the plantar surface of the hind paw and the time taken to withdraw from the heat stimulus is recorded (44). For cold measurement, a dry ice method was used to deliver the cold stimulus to the glass underneath the hind paw and the latency was recorded (45).

#### SHIM incapacitance meter to measure static limb use

Static weight bearing was measured using a SHIM incapacitance meter (IC Meter) connected to a system 8000 micro-measurements tool (15). Data were collected from mice as described previously (15). Data were retained if the animal stood still in relaxed position, without noticeably shifting weight, lifting or offloading a limb, or turning the head. Each animal was tested until five appropriate trials were obtained. Hind limb distribution of weight was recorded using the strain smart software, and transferred to data files. Following data collection, mice were placed back into their home cage. Weight bearing on the limb of interest (ipsilateral [right] or contralateral [left]) was expressed as a percentage of the total weight born by the hindlimbs, as per the equation:

([weight placed on ipsilateral or contralateral]/ [total weight placed on both hindlimbs])*100 SHIM data were also expressed as symmetry indices (SI) according to the equation:

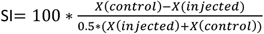

### Assessment of the hyperalgesic effects of artemin

Naïve male C57BL6 mice received hind paw injections of artemin to determine whether artemin can induce hypersensitivity. A 200 ng injection of Artemin (R&D Biosystems; Cat. 1085-AR/CF) was delivered in 10 μl of sterile saline using an insulin syringe to the subcutaneous layer of the left hind paw while mice were conscious and restrained. Control mice received 10 μl of sterile saline. Mice were assessed for weight-bearing and for mechanical, heat, and cold sensitivity. A single behavioral test was performed following each injection to avoid overstressing them with multiple behavioral assays, as they were being tested at 1, 2, 4, 6, and 24 hours post-injection. Mice had a minimum of one week of washout between hind paw injections.

### Immunohistochemistry

Lumbar DRGs were isolated from naïve, control, and MIA-injected mice, sectioned on a cryostat at 12 μm thickness for double immunofluorescence labeling (46). Primary antibodies were diluted (1:500 each) in 5% blocking solution. The primary antibodies used were Tuj1 (Abcam; Cat: ab7751), a neuronal marker, TRPV1 (Santa Cruz, Cat: Sc398417) and GFRα3 (Neuromics, Cat: GT15123,). Alexa Fluor conjugated secondary antibodies (Invitrogen; 488, Cat: A20181; 546, Cat: A20183) were applied in 2% blocking solution for 1 hour. Sections were washed, dried, and finalized with mounting media contained DAPI and imaged using a Leica microscope. Counting was performed on the images by an individual who was blinded to group allocation. DRG images were analyzed for the total number of neurons (Tuj1-positive), and for the total number of neurons expressing GFRα3 (Tuj1- and GFRα3-positive). Ipsilateral (MIA) and contralateral DRG were analyzed and compared.

### Assessment of the analgesic effect of a systemic anti-artemin monoclonal antibody

The analgesic effects of blocking artemin were determined using mice with MIA-induced OA pain. Following baseline behavioral tests (heat, mechanical, and cold sensitivity testing) mice had OA induced using intra-articular MIA (as described above). Behavioral tests were performed every week up to day 28. Following day 28, mice received a 100 μl IP injection of either PBS or 25 μg of anti-artemin monoclonal antibody (R&D, cat: MAB10851-500) dissolved in PBS. Allocation to treatment was randomized, and testing was performed by an individual blind to treatment allocation (LM). Mice were tested for sensitivity to heat, cold, and mechanical stimuli at 2, 4, 6, and 24 hours post intraperitoneal injection.

### Statistical Analysis

All behavioral data were collected by a researcher blind to the groups. Data were tested for normalcy with the Shapiro-Wilk test. Two-way ANOVA and t-tests with corrections for multiple comparisons were performed using GraphPad Prism. For multiple t-tests, statistical significance determined using the Holm-Sidak method, using a corrected p-value of 0.05.

## Data availability

All data will be shared on reasonable request to.

## Authors Contribution

SKM, BDXL conceptualize the study and designed experiments, LM performed the experiments and analyzed the data with SKM and BDXL. SKM and BDXL wrote the manuscript and all authors approved it.

## Funding and additional information

Funding for this work was provided by donations to the Translational Research in Pain (TRiP) Program from individuals interested in supporting an understanding of the mechanisms of osteoarthritis pain.

---

**Supplementary Figure 1.**
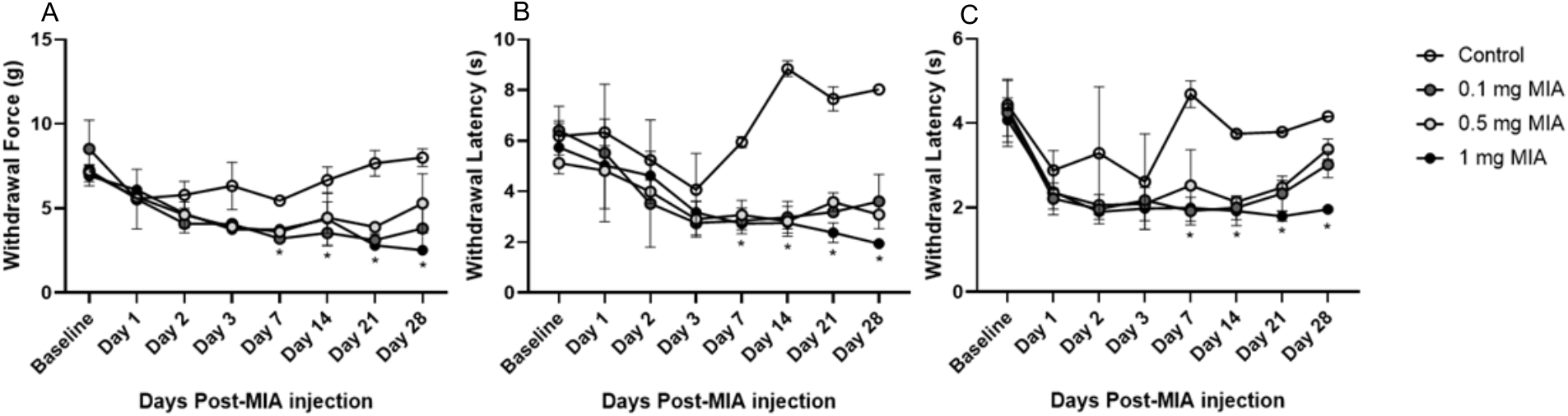
Hypersensitivity in response to multiple doses of MIA. Mice were tested for hypersensitivity to mechanical (A), heat (B), and cold (C) stimuli using the Von Frey, Hargreaves, and cold assays through 28 days. Control (n=2), 0.1 mg MIA (n=2), 0.5 mg MIA (n=2), and 1 mg MIA (n=2). Data are represented as mean ± SD. There was significant difference between control mice and 1 mg MIA at day 28 for all three behavioral assays (two-tailed t-test assuming unequal variances; Von Frey p<0.05; Hargreaves p<0.05; Cold p<0.01). ANOVA with repeated measures for day 7 through day 28 found that all MIA doses were significantly different from control mice after MIA induction for all three assays (p<0.05).

**Supplementary Figure 2.**
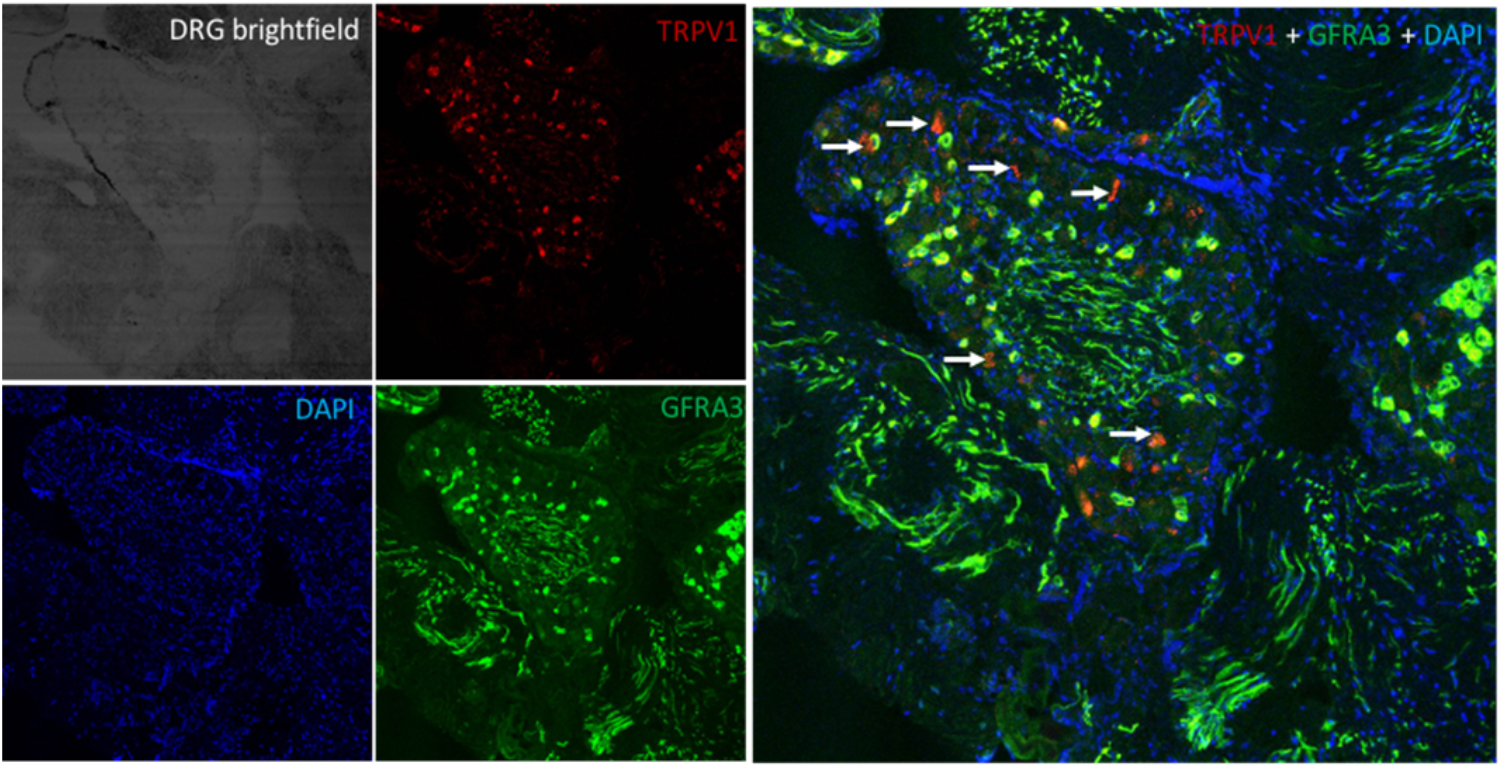
Co-expression of GFRα3 and TRPV1 receptors. A representative image from. double IHC reveals a co-expression of GFRα3 and TRPV1 receptors in normal mouse DRG. Three independent mice were shown similar co-expression profile.

